# Virus-human protein-protein interactions predict viral phenotypes

**DOI:** 10.64898/2026.06.12.732009

**Authors:** Zhiyuan Zhang, Yang Feng, Xingyi Ge, Xiangxian Meng, Yousong Peng

## Abstract

Viral phenotypes such as host and tissue tropism are critical determinants of viral infection and transmission. Inferring viral phenotypes presents unique challenges compared to cellular organisms, as viruses rely entirely on host machinery for replication and survival. Current methods for predicting viral phenotypes mainly rely on viral genomic data, often overlooking host-related information. Here, we evaluated the utility of predicted virus-human protein-protein interactions (PPIs) in inferring diverse viral phenotypes using machine-learning algorithms. For predicting human infectivity, a PPI-based machine learning model outperformed both virus genomic and protein sequence-based models that used large language model embeddings. It also surpassed previous methods that incorporated both viral and host genomic data. The human proteins identified by the model were significantly enriched in functions related to viral infection and immune response. In predicting various phenotypes of human RNA viruses, PPI-based models performed better than virus sequence-based models in forecasting virulence, human transmissibility and transmission routes, while showing comparable performance to genomic sequence-based models in predicting tissue tropism. Finally, we demonstrated that a PPI-based model could distinguish high-risk HPV genotypes from low-risk ones. Proteins associated with high-risk HPV were involved in apoptosis and immune regulation, whereas those linked to low-risk HPV were enriched in telomere maintenance and DNA repair. Collectively, this study is the first to demonstrate the value of predicted virus-human PPIs in inferring viral phenotypes, thereby enhancing our understanding of the molecular mechanisms underlying these phenotypes. It also provides effective tools for risk assessment of emerging viruses, contributing to improved pandemic preparedness.

## Introduction

Viruses are ubiquitous in the globe and have caused substantial damages to humans. They can cause numerous viral diseases such as viral hepatitis and tumor[1]. They also cause lots of epidemics or pandemics in human history and lead to numerous human deaths. For example, the most recent pandemic by the Severe Acute Respiratory Syndrome coronavirus 2 (SARS-CoV-2) has caused more than 7 million deaths[2]. Most of these viruses are zoonotic and cross-species to infect humans. Timely surveillance and risk evaluation of zoonotic viruses are key for prevention and control of newly emerging viruses.

Determining the viral phenotypic traits such as human infectivity, virulence, transmissibility, transmission route, pathogenicity, tissue tropism and carcinogenicity is a pre-requirement for accurate risk evaluation of zoonotic viruses and precise medicine of viral diseases. Experimental studies of viral phenotypes primarily rely on *in vitro* and *in vivo* models. The *in vitro* models included two-dimensional (2D) and three-dimensional (3D) cell culture models. The 2D models such as Vero cell and A549 cells have been commonly used in isolation and characterization of viruses. The 3D models such as explants and organoids more closely mimic human physiology compared to the 2D models. For instance, Porotto et al. used the lung organoids to investigate the impact of respiratory virus infections on the developing lung of children[3]. I*n vitro* models cannot fully recapitulate the dynamic physiological environment of living organism, making *in vivo* animal models indispensable. The mouse is the most widely used animal model in virus phenotype studies. For example, Xiang et al. investigated the pathogenicity of Echovirus 18 using the mouse model[4]. Although experimental approaches are the mainstream methods in virus characterization studies, they remain time-consuming, labor-intensive, and costly, and require stringent biosafety conditions due to the inherent pathogenic risk.

Computational methods have emerged as valuable tools for predicting viral phenotypes and can generally be classified into two categories: sequence-based and trait-based approaches. Lots of sequence-based methods have been developed for virus phenotype prediction. For example, Davis et al. predicted the human pathogenicity of novel coronaviruses using a Logistic Regression model trained on k-mer frequencies of viral genomic sequences[5]; Zhang et al. and Alshayeji et al. predicted the human-infecting viromes using sequence features based on machine-learning methods[6,7]; Li et al. inferred multiple phenotypes of influenza A viruses such as human adaptation, pathogenicity, and drug-resistance based on genomic sequences[8]. Most of these methods only considered the virus sequences, ignoring the virus-host interactions, and are virus-specific, which limited their applications in newly emerging viruses. The trait-based methods predict virus phenotypes using prior knowledge of viruses. For example, Brierley et al. predicted virulence of human RNA viruses based on tissue tropism and transmission ecology of viruses[9]. Unfortunately, there is little prior knowledge for most emerging viruses.

Viruses rely on host cells for survival. For infections of host cell, viruses have to enter the host cell, inhibit the host cell immunity and then use the host cell machine for replication and generation of progeny virion. All these processes rely on close virus-host protein-protein interactions (PPIs). For instance, the interaction between SARS-CoV-2 spike protein and the human ACE2 receptor mediates viral entry into host cells and determines host susceptibility[10], while influenza A virus nonstructural protein 1 (NS1) targets the human TRIM25 (a ubiquitin ligase) to suppress innate immune signaling and promote viral replication and spread[11]. Thus, virus-host PPIs are key determinants in viral infections and pathogenicity. In our previous work, we developed vhPPIpred, a machine learning framework that integrates sequence embeddings and evolutionary information with network topology and viral molecular mimicry to predict virus-human protein-protein interactions (PPIs)[12]. While this previous work focused on the prediction and characterization of virus-human PPIs themselves, whether these interaction profiles can be further leverage to infer viral phenotypes remains unexplored. Here, we addressed this question by developing a novel computational framework that leverages predicted virus-human PPIs to predict five kinds of viral phenotypes: human infectivity, virulence, transmissibility, transmission route, and tissue tropism. The framework contains two main components: (1) large-scale prediction of virus-human PPIs, and (2) machine learning modeling of virus phenotypes with the predicted PPIs. Comparative experiments showed that PPI-based models consistently outperformed genome- or proteome-based models across all phenotype prediction tasks. Our study therefore extends virus-human PPI prediction from identifying molecular interactions to leveraging interaction profiles for systematic viral phenotype inference, providing new insights into viral pathogenesis and facilitates the risk evaluation of newly emerging viruses.

## Results

### Framework for predicting viral phenotypes based on virus-human PPIs

The ability to anticipate the public health risk posed by viruses critically depends on understanding their biological traits. However, for most newly identified or poorly characterized viruses, these phenotypes remain unknown. Virus-host interactions, particularly protein-protein interactions (PPIs) are key determinants of viral phenotypes. Therefore, we proposed a computational framework for predicting viral phenotypes based on virus-human PPIs. For this, we firstly manually compiled a comprehensive dataset of virus-human PPIs that contains 16,314 pairs of experimentally determined physical virus-human PPIs (Table S1). Unfortunately, the dataset is far from complete as it only covered 120 virus species and 80% viruses had less than 80 PPIs (Figure 1A and Table S1). Besides, it is biased towards some viruses such as SARS-CoV and influenza viruses that contain more than 50% of these PPIs (Figure 1A). To overcome these limitations, we developed vhPPIpred, a machine learning-based prediction method that not only incorporated sequence embedding and evolutionary information but also leveraged network topology and viral molecular mimicry of human PPIs, to predict virus-human PPIs based on the experimentally-determined physical virus-human PPIs [12]. vhPPIpred outperformed nine state-of-the-art methods (including AlphaFold3) on both our benchmark dataset and three independent datasets. Thus, the high-confidence virus-human PPIs predicted by vhPPIpred were taken as features in machine-learning modeling of five key viral phenotypes including human infectivity (HI), virulence (VL), human transmissibility (HT), transmission route (TR), and tissue tropism (TT) (Figure 1B).

**Figure 1.**
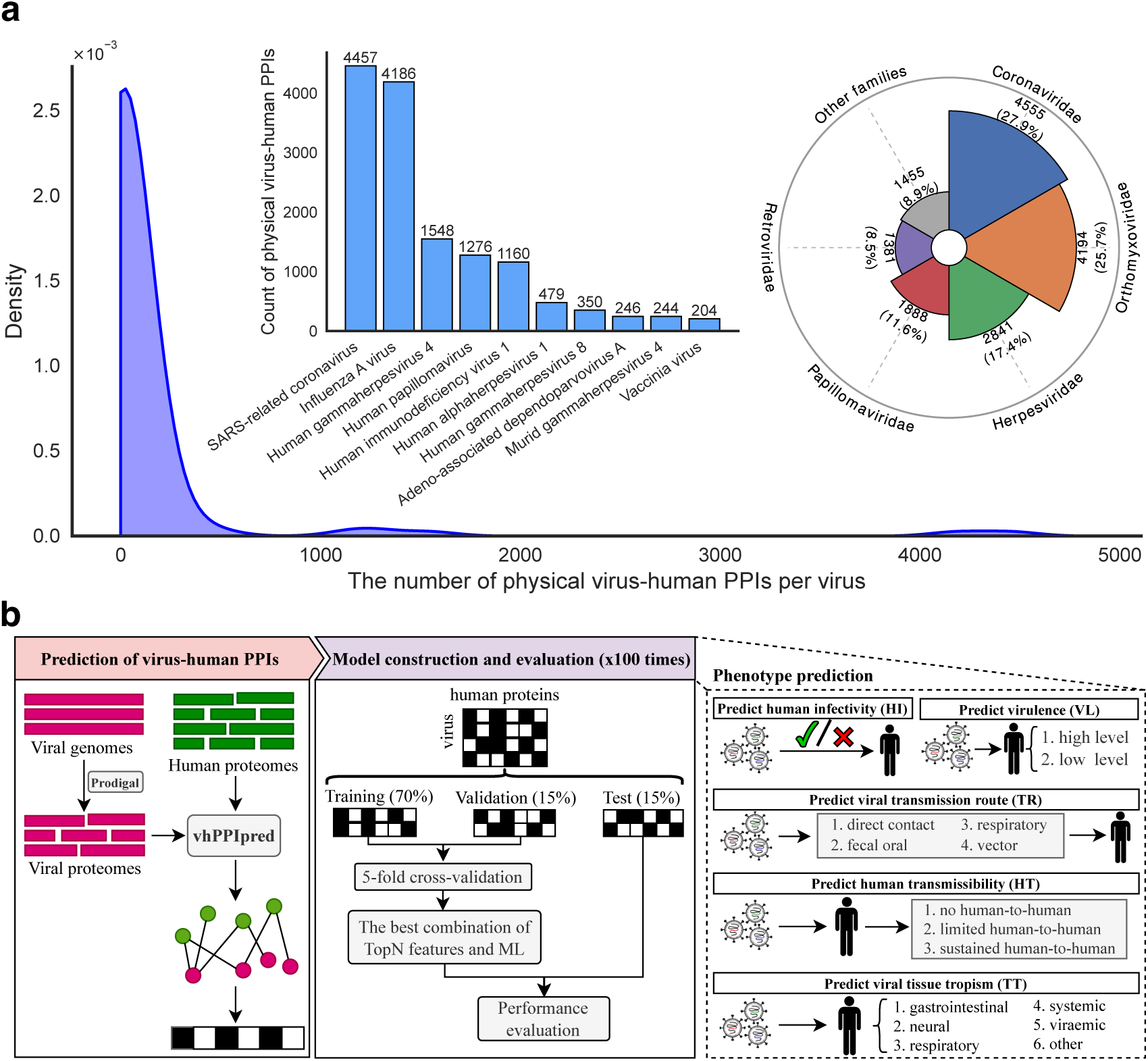
Schematic overview of the PPI-based computational framework for viral phenotype prediction. **A**. Distribution of experimentally determined physical virus-human PPIs. The top 10 virus species and top 5 families of all PPIs were listed in the bar and pie chart, respectively. **B**. Framework for predicting viral phenotypes with predicted virus-human PPIs.

### Prediction of human infectivity of viruses

Human infectivity (HI) determines whether a virus can infect human. Series of approaches have been proposed for human infectivity prediction. For example, Zhang et al. discriminated human-infecting viruses from other viruses with K-Nearest Neighbors (KNN) algorithm based on k-mer frequencies of viral genomic sequences[6]. Mollentze et al. identified human viruses using eXtreme Gradient Boosting (XGBoost) algorithm based on host signatures encoded in viral genomes[13]. However, the number of human-infecting viruses used in these studies was small compared to that estimated by the Global Virome Project[14]. Thus, we firstly constructed a comprehensive dataset of human-infecting viruses from literatures and multiple databases such as Virus-Host Database and VIRION (Figure 2A). The dataset contains 1,469 human-infecting viruses from 49 viral families which are more than twice that used in previous studies (Figure 2A and Table S2). The human-infecting viruses were categorized into high, middle and low-confidence groups comprising 331, 346, and 792 species, respectively, according to the evidence of human infections (Table S2) (see Materials and Methods). Only 677 viruses from the high and middle-confidence groups were used in the modeling for reliable prediction of human-infections and served as positive samples. They were distributed in 48 families (Figure 2B and Table S2), among which *Papillomaviridae*, *Anelloviridae*, and *Flaviviridae* contained the greatest number of human-infecting viruses. A similar number of high-confidence non-human-infecting viruses (n=564) were compiled from Mollentze’s study and served as negative samples in the modeling (Table S3).

**Figure 2.**
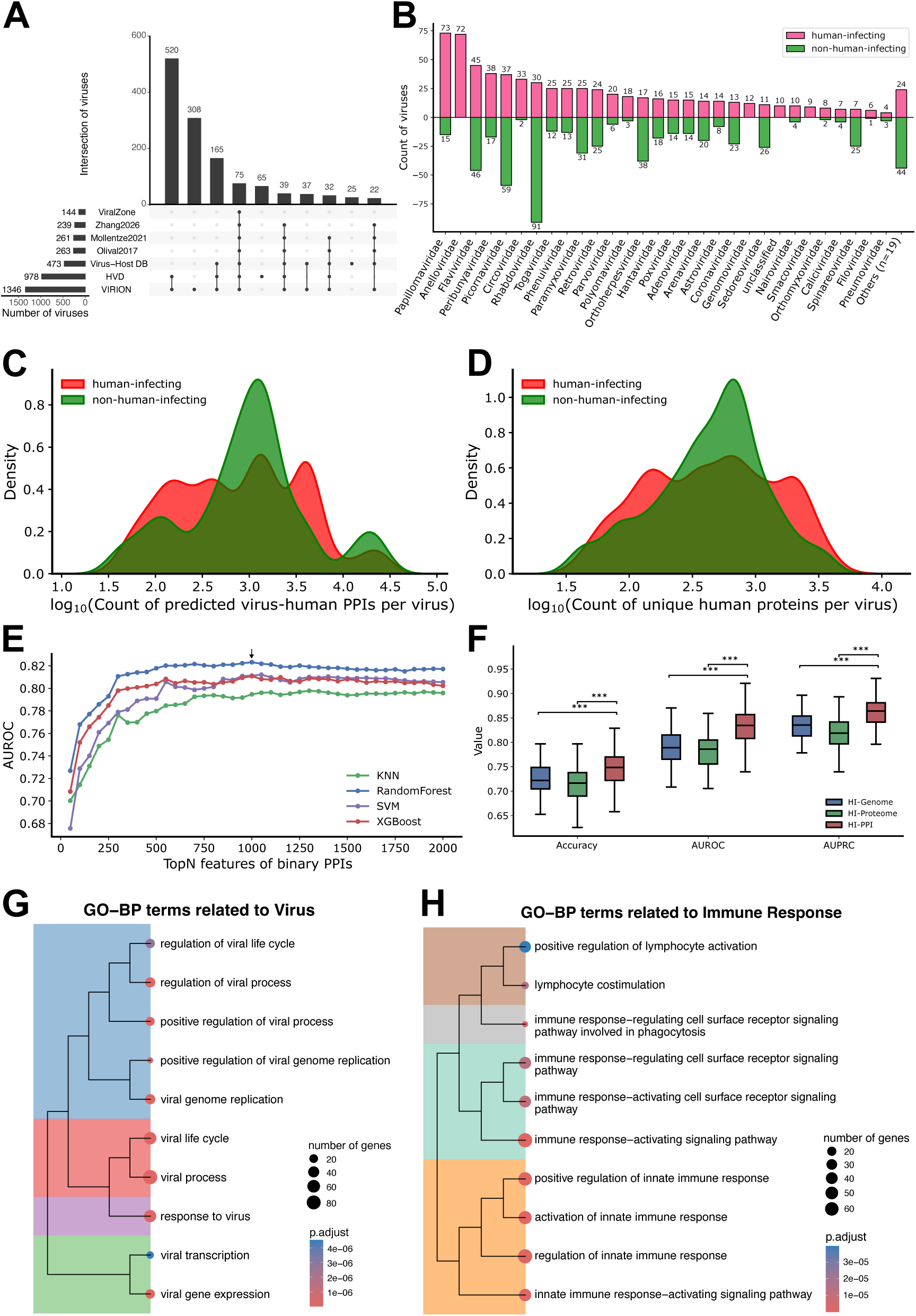
Prediction of viral human infectivity. **A.** Data sources of all human-infecting viruses (including high-, middle-, and low-confidence viruses). **B.** Distribution of human-infecting and non-human-infecting viruses across viral families. **C.** Distribution of predicted virus-human PPIs between human-infecting and non-human-infecting viruses. **D.** Distribution of unique human proteins predicted to interact with viruses between human-infecting and non-human-infecting viruses. **E.** Validation AUROC across different combinations of machine learning algorithms and the top N (N=50, 100, …, 2000) binary PPI features during feature selection. **F.** Comparison of accuracy, AUROC, and AUPRC among the best genome-, proteome-, and PPI-based models on the test sets. **G-H**. Functional clusters of enriched Gene Ontology biological process (GO:BP) terms related to viral replication and host-virus interaction (**G**), and those related to host immune regulation and antiviral defense (**H**) among the top 450 human proteins used in the HI-PPI model.

For both human-infecting and non-human-infecting viruses, high-confidence PPIs between virus and humans were predicted using vhPPIpred (see Materials and Methods). The number of virus-human PPIs predicted varied much by viruses, ranging from fewer than 100 to over 10,000 (Figure 2C). The median number of predicted PPIs was 943 and 1,076 for human-infecting and non-human-infecting virus, respectively. Similarly, the number of human proteins that were predicted to interact with viruses also varied much across viruses, ranging from 35 to 5,538 (Figure 2D). No significant difference was observed in the number of predicted virus-human PPIs (p-value=0.055 in the two-sided t test, Figure S1A), or number of human proteins that were predicted to interact with viral proteins (p-value=0.604 in the two-sided t test, Figure S1B), between human-infecting and non-human-infecting viruses. Further analysis showed that the number of predicted virus-human PPIs was strongly positively correlated with the genome size (Pearson r=0.919, Figure S1C) and the number of viral proteins (Pearson r = 0.899, Figure S1D). These results suggest that the high number of predicted human interactors for some viruses may be partly explained by their larger genomes which encode a larger repertoire of viral proteins.

All human proteins that were predicted to interact with viral proteins were taken as features in the modeling. Specifically, for each virus, it was represented as a vector of virus-interacting human proteins. Then, feature and algorithm selections were conducted to obtain the optimal model performances. All virus samples (including both human-infecting and non-human-infecting viruses) were divided into training, validation, and test sets, with the ratio of 70%, 15%, and 15%, respectively. The training and validation sets were used for feature-algorithm selection and model construction using 5-fold cross-validation, while test set was used for model evaluation. All these processes were repeated 100 times, resulting in average metrics. The virus-interacting human proteins (features) were ranked by the importance that was measured with SHapley Additive exPlanations (SHAP) values. As shown in Figure 2E, the AUROC of models increased with the number of features used that were ranked by importance, then they began to decline after a certain number of features. The model performance (AUROC) varied among four machine-learning algorithms including Random Forest (RF), K-Nearest Neighbors (KNN), Support Vector Machine (SVM) and eXtreme Gradient Boosting (XGBoost). The RF and XGBoost consistently outperformed other two algorithms. The model with the combination of RF algorithm and top 1000 features achieved the best performance in the cross-validations, with average AUROC of 0.823. It was selected to predict human infectivity in the study and named as HI-PPI. Evaluation of the model on the test sets showed that it achieved an average AUROC of 0.830 (Table S4). More specifically, the average accuracy and AUPRC are 0.749 and 0.861, respectively (Figure 2F and Table S4).

For comparison, we also utilized viral genomic and proteomic sequence features to model human infectivity using the same procedure and the same dataset as the PPI-based modeling. For viral genome sequences, two kinds of features were used: one was k-mer frequencies (k=3,4,5) of genomic sequences, and the other was sequence embeddings by DNABERT-2 (a foundation model trained on large-scale multi-species genomes)[15]. Similarly, viral proteomes were also represented with k-mer frequencies (k=1,2,3) and sequence embeddings by ESM-2 (a transformer-based language model trained on protein sequences of UniRef50)[16]. Models with the combination of 3-mer frequencies and RF algorithm and with the combination of ESM-2 embeddings and RF algorithm were selected as the optimal model, respectively, for the genome-based and proteome-based models (Figure S2), and were named as HI-Genome and HI-Proteome, respectively. Both models were subsequently tested on the test sets. As shown in Figure 2F, the AUROC of both models were significantly lower than that of HI-PPI (p-value < 1 × 10^−3^ in the two-sided t test). Other metrics including accuracy, precision, recall, F1-score, AUPRC of both models were also 0.02-0.04 lower than those of HI-PPI (Table S4).

We also compared our PPI-based method with Mollentze’s method on their dataset which contained 261 human-infecting and 600 non-human-infecting viruses. The results showed that the PPI-based method achieved an average AUROC of 0.842, which was significantly higher than that of Mollentze’s model (0.842 vs. 0.737, p-value = 5.76 × 10^−46^ in the two-sided t test, Figure S3B).

Finally, we conducted interpretability analysis for PPI-based methods by functional enrichment analysis on the top 1000 human proteins used in HI-PPI. Interestingly, several virus-related biological process including viral gene replication, expression, host response to viruses, and regulation of these processes were significantly enriched for these human proteins, suggesting they were involved in viral infection process (Figure 2G). Similarly, several immune-related biological processes including regulation of lymphocyte activation, regulation of innate immune response, and immune response-regulation signaling pathway were also enriched for these human proteins (Figure 2H). Among these immune-related human proteins, most of them were associated with terms like “activation of innate immune response”, and “negative regulation of immune response”. This suggests that immune-related protein were also important determinants of viral human infectivity.

### Prediction of diverse viral phenotypes for RNA viruses

Besides human infectivity, other viral traits including virulence (VL), human transmissibility (HT), transmission route (TR), and tissue tropism (TT), are also critical for evaluating their zoonotic potential and pandemic risk. Thus, we further evaluated the effectiveness of virus-human PPIs in predicting these viral traits of RNA viruses using the same procedure as the human infectivity modeling. All viruses and related viral traits were curated from Brierley’s study[9]. They were divided into training, validation, and test sets as described above in modeling of each viral trait.

For virulence (VL) prediction, a total of 214 RNA viruses were assigned into either the low-level virulence group, which included viruses causing non-severe human diseases (n=155), or the high-level virulence group, which included viruses associated with severe human diseases (n=59) (Figure 3A, Table S5). During feature and algorithm selection, the PPI-based model using an SVM classifier and the top 250 virus-interacting human proteins (named as VL-PPI) achieved the best cross-validation performance, with an AUROC of 0.831 (Figure S4A). To benchmark this model against sequence-derived predictors, genome- and proteome-based models were also constructed. The models with the combination of SVM algorithm and DNABERT-2-derived genome representations (named as VL-Genome), and with combination of KNN algorithm and 3-mer frequencies (named as VL-Proteome) were selected for the genome and proteome-based model, respectively. In cross-validation, both VL-Genome (0.675) and VL-Proteome (0.678) had a lower AUROC than the VL-PPI in the cross-validations (Figure S4A, E, and I). Further evaluation on the test sets showed that VL-PPI had a comparable accuracy to VL-Genome and VL-Proteome (0.799 vs. 0.795, and 0.789) (Figure 3J), but achieved consistently better discriminatory performance, including AUROC (0.844 vs 0.763 and 0.781) and AUPRC (0.708 vs 0.636 and 0.607) (Figure 3B&F and Table S6).

**Figure 3.**
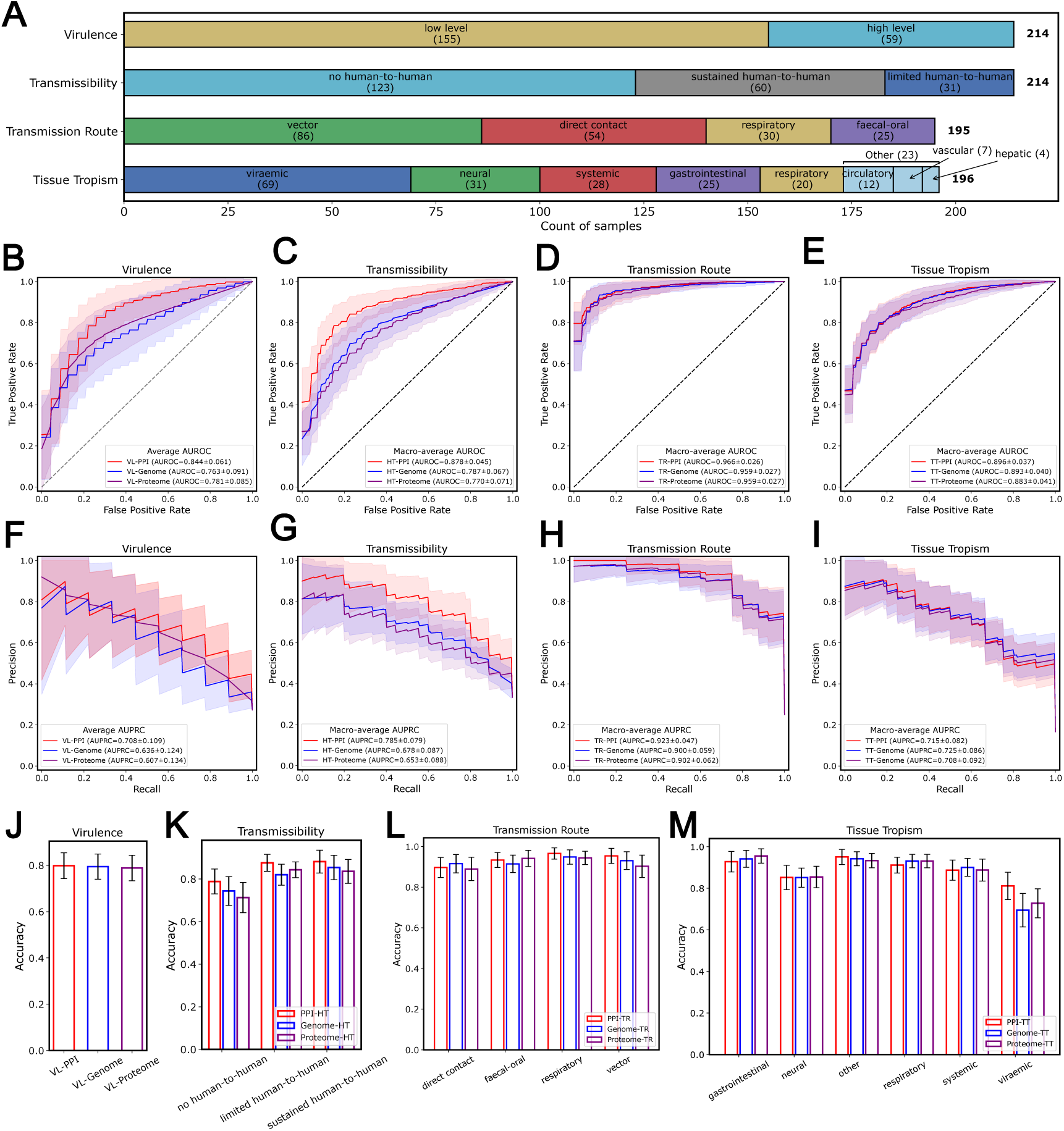
Comparison of PPI- and sequence-based models for predicting human transmissibility, transmission route, and tissue tropism of RNA viruses. **A.** Components of RNA viruses used for predicting four phenotypes. B-E. Comparison of AUROC (or macro-AUROC) between PPI- and sequence-based models for virulence (B), transmissibility (C), transmission route (D), and tissue tropism (E) on the test sets. F-I. Comparison of AUPRC (or macro-AUPRC) of models between PPI- and sequence-based models for virulence (F), transmissibility (G), transmission route (**H**), and tissue tropism (**I**) on the test sets. **J-M.** Accuracy of virulence prediction (**J**) and models across different categories within transmissibility (**K**), transmission route (**L**), and tissue tropism (**M**).

For human transmissibility (HT) prediction, the same set of 214 RNA viruses were classified into three levels according their capacity for human-to-human transmission: low, defined as no human-to-human transmission (n = 123); middle, defined as limited human-to-human transmission (n = 31); and high, defined as sustained human-to-human transmission (n = 60) (Figure 3A, Table S5). Feature and algorithm selection showed that the PPI-based model with the combination of SVM algorithm and top 200 virus-interacting human proteins (named as HT-PPI) achieved the best performance in the cross-validations, with the macro-AUROC of 0.871 (Figure S4B). For comparison, sequence-based models were built as mentioned above. The models with the combination of KNN algorithm and 4-mer frequencies (named as HT-Genome), and with the combination of RF algorithm and amino acid frequencies (named as HT-Proteome) were selected for the genome and proteome-based model, respectively (Figure S4F&J). Both HT-Genome (0.766) and HT-Proteome (0.760) had a lower macro-AUROC than the HT-PPI in the cross-validations. Further evaluation on the test sets showed that HT-PPI significantly outperformed both HT-Genome and HT-Proteome on nearly all metrics, such as macro-AUROC (0.878 vs 0.787 and 0.770) and macro-AUPRC (0.785 vs 0.678 and 0.653) (Figure 3C&G). Class-wise evaluation further showed that HT-PPI performed particularly well for viruses with middle and high transmissibility, reaching accuracies of 0.875 and 0.881, respectively, whereas its accuracy for low-transmissibility viruses was relatively lower at 0.788. Nevertheless, the performance of HT-PPI across all transmissibility levels remained significantly higher than that of both sequence-based models (Figure 3K and Table S6).

For transmission route (TR) prediction, 195 RNA viruses with recorded TR were divided into four categories: vector-borne (n = 86), direct contact (n = 54), respiratory (n = 30), and faecal-oral (n = 25) (Figure 3A, Table S5). Feature and algorithm selection showed that the PPI-based model with the combination of SVM algorithm and top 350 virus-interacting human proteins (named as TR-PPI) achieved the best performance in the cross-validations, with the macro-AUROC of 0.969 (Figure S4C), which slightly surpassed the best genome-based model with the combination of SVM algorithm and 4-mer frequency (named as TR-Genome, macro-AUROC=0.957) and the best proteome-based model with the combination of RF algorithm and ESM-2 embedding (named as TR-Proteome, macro-AUROC=0.956) (Figure S4G&K). Evaluation on the test set confirmed the advantage of PPI-based models, as TR-PPI exhibited higher macro-AUROC (0.966 vs. 0.959 and 0.959) and macro-AUPRC (0.923 vs. 0.900 and 0.902) than both TR-Genome and TR-Proteome (Figure 3D&H). Analysis of model performances by TR showed that TR-PPI can predict viruses with each kind of TR accurately, with accuracies greater than 0.85 for all TRs. TR-PPI also outperformed TR-Genome and TR-Proteome on all TRs except the direct contact (Figure 3L and Table S6).

For tissue tropism (TT) prediction, 196 RNA viruses with recorded tissue tropism were categorized into six groups: viraemic (n = 69), neural (n = 31), systemic (n = 28), gastrointestinal (n = 25), respiratory (n = 20), and other (including circulatory (n = 12), vascular (n = 7), and hepatic (n = 4)) (Figure 3A). Feature and algorithm selection showed that the PPI-based model with the combination of SVM algorithm and top 1250 virus-interacting human proteins (named as TT-PPI) achieved the best performance in the cross-validations, with the macro-AUROC of 0.886 (Figure S4D), which slightly surpassed the best genome-based model with the combination of RF algorithm and ENABERT2 encoding (named as TT-Genome, macro-AUROC=0.885) (Figure S4H) and the best proteome-based model with the combination of RF algorithm and ESM-2 embedding (named as TT-Proteome, macro-AUROC=0.878) (Figure S4L). Evaluation on the test set showed that the TT-PPI performed similarly as the TT-Genome, with close macro-AUROC (0.896 vs. 0.893) and macro-AUPRC (0.715 vs. 0.725) (Figure 3E&I). Both TT-PPI and TT-Genome outperformed the TT-Proteome. Analysis of model performances by TT showed TT-PPI achieved high accuracies in predicting virus with tropism towards gastrointestinal, respiratory and other tissues (accuracy > 0.90), while had relative low accuracies in viraemic and neural tissues (Figure 3M and Table S6). It slightly outperformed sequence-based models in predicting viruses with tropism towards neural, viraemic, and other tissues (Table S6).

Collectively, these results demonstrated that virus-human PPIs could be used to predict multiple phenotypes of RNA virus with high accuracy and improve the prediction performance of machine-learning models compared to those based on genome or proteome sequence features (including the large language model embeddings). The superior and more consistent performance of PPI-based models across diverse tasks suggested that virus-human interaction patterns capture biologically meaningful determinants of viral behaviors, reflecting important roles of virus-human PPIs in viral infection and transmission. This emphasizes the value of integrating host information to better assess the zoonotic and pandemic potential of emerging viruses.

### Phenotype inference and risk evaluation of potentially human-infecting viruses using PPI-based models

As mentioned above, a total of 792 low-confidence of human-infecting viruses were curated, including 119 DNA viruses, 636 RNA viruses, and 37 unclassified viruses (Table S2). The human infectivity of these potentially human-infecting viruses was predicted using the HI-PPI model. As shown in Figure S5, a total of 450 viruses were predicted to infect humans, with the score ranging from 0.510 to 0.920 (Figure S5). They were classified into 32 viral families, among which 29 viral families contained known human-infecting viruses (Figure 4A, Table S8). The family of *Picornaviridae* contained the largest number of potentially human-infecting viruses (160) and the ratio of known human-infecting viruses is 23%. Families like *Caliciviridae* and *Astroviridae* have over 10 potentially human-infecting viruses and the known human-infecting viruses of them account for more than 30%. These results on one hand demonstrated the reliability of the potentially human-infecting viruses, and on the other hand validated our PPI-based method in predicting viral human infectivity.

**Figure 4.**
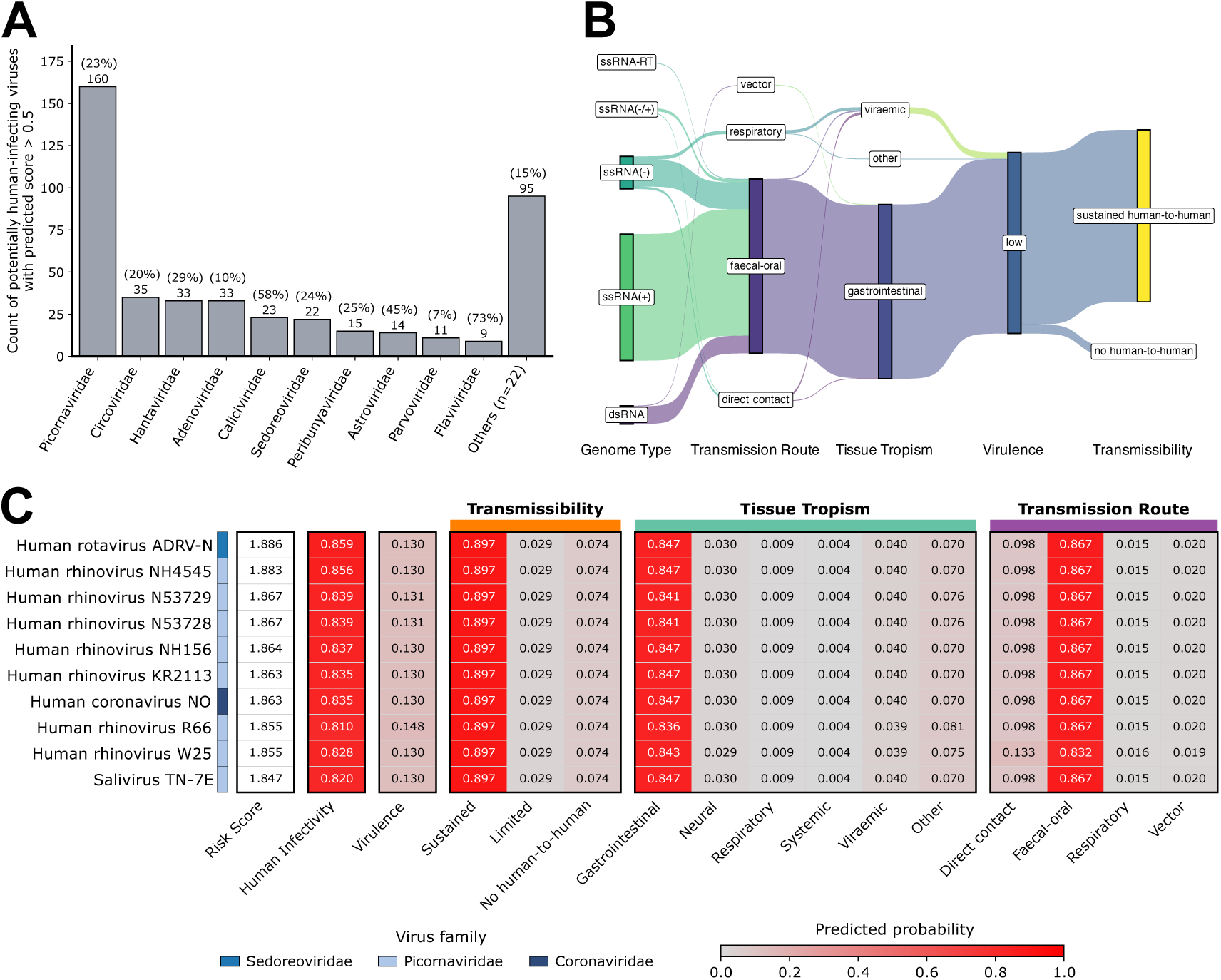
Phenotype predictions of potentially human-infecting viruses using PPI-based models. **A.** Distribution of potentially human-infecting viruses across viral families. Bar heights indicate the number of viruses predicted to infect humans in each viral family. Percentage above bars represent the proportion of known human-infecting viruses (high- and middle-confidence levels) relative to all viruses classified by the International Committee on Taxonomy of Viruses (ICTV) within the corresponding viral family. **B.** Predicted transmission route, tissue tropism, virulence, and transmissibility of potentially human-infecting RNA viruses. **C.** Top10 human-infecting RNA viruses of concern ranked by risk score, which was the summary of predicted scores of human infectivity, virulence and sustained human-to-human transmissibility.

For further risk evaluation of these potentially human-infecting viruses, we applied the models developed above (VL-PPI, HT-PPI, HR-PPI and TT-PPI) to predict their virulence, human transmissibility, transmission route, and tissue tropism. Since all these models were trained on RNA viruses, only 319 potentially human-infecting RNA viruses were used here. Most of these RNA viruses possessed the positive-sense single-stranded RNA (ssRNA(+)) genome, followed by those with a negative-sense single-stranded (ssRNA(-)) genome (Figure 4B). Interestingly, among these viruses, 96.2% (307/319) were predicted to be transmitted via the faecal-oral route and to predominantly target the gastrointestinal tract. All were predicted to have low virulence, and 95% (303/319) were predicted to exhibit sustained human-to-human transmissibility. In contrast, only a small proportion were predicted to be transmitted via the respiratory route (1.9%, 6/319) or exhibit viraemic tropism (3.5%, 11/319) (Figure 4B, Table S9). Subsequently, these viruses were ranked by the overall risk score (sum of the scores of human infectivity, virulence, sustained human-to-human transmission) and the top 10 viruses were shown for illustration. As shown in Figure 4C, the top 10 ranked viruses belonged to three viral families and were consistently predicted to exhibit sustained human-to-human transmissibility, gastrointestinal tropism, low virulence, and predominantly faecal-oral transmission. Notably, several top-ranked viruses, although classified as low-confidence based on our predefined evidence criteria, belonged to viral groups with well-established relevance to human diseases, including human rotaviruses associated with gastroenteritis[17], as well as human rhinoviruses[18] and coronaviruses[19]. These observations demonstrate consistency between our predictions and existing evidence of human infection in the corresponding viral groups.

### Classification of high and low-risk HPVs based on virus-human PPIs

We further investigated the ability of PPI-based method in inferring phenotypes at the virus sub-species or genotype level. Human Papillomavirus (HPV) is a common human pathogen and contained more than 150 genotypes, of which approximately 40 genotypes infected the anogenital tract such as HPV-16 and HPV-18[20]. It can be classified into two types according to its ability of causing cancers: high-risk HPVs (HR-HPVs) that can lead to multiple cancers such as the cervical cancer, and low-risk HPVs (LR-HPVs) that are primarily related to benign warts. A total of 38 HPV genotypes with known pathogenicity including 12 HR-HPVs and 26 LR-HPVs were collected from Lasso’s study (Figure 5A)[21]. PPIs between each HPV genotype and humans were predicted and were used in classifying HR-HPV and LR-HPV with machine-learning algorithms as mentioned above. The top 30 most important virus-interacting human proteins and the random forest algorithm were selected via the cross-validations on the training set (Figure 5B). Evaluation of the model on the test set suggests its outstanding ability in distinguishing HR-HPVs and LR-HPVs, with an average AUROC of 0.903, AUPRC of 0.866, and accuracy of 0.859 (Figure 5C). These results demonstrated that virus-human PPIs can be used to effectively classify HR-HPVs and LR-HPVs. Then, the interactions between HPVs and the top 30 virus-interacting human proteins used in the modeling were visualized to explore the role of virus-human PPIs in determining virus pathogenicity. As shown in Figure 5D, the genes encoding these proteins could be clearly divided into two groups based on their interactions with HPVs: HR-HPV-related group that mainly interact with HR-HPV (16 genes, e.g., CHCHD10, ACIN1, SSBP1, FAM209A, TNFRSF1A) and LR-HPV-related group that mainly interact with LR-HPV (14 genes, e.g., GRIPAP1, TERF2, CALU, EPB41L1, STX17). Reactome enrichment analysis showed that HR-HPV-related genes were clustered in TNF/NF-κB-related pathways involved in apoptosis and immunity, supporting immune evasion and tumorigenesis (Figure 5E)[22]. In contrast, LR-HPV-related genes were clustered in pathways related to telomere maintenance and DNA repair, which might contribute to genome stability without clear carcinogenic potential (Figure 5F). These results were consistent with the distinct pathogenic characteristics of HR- and LR-HPVs.

**Figure 5.**
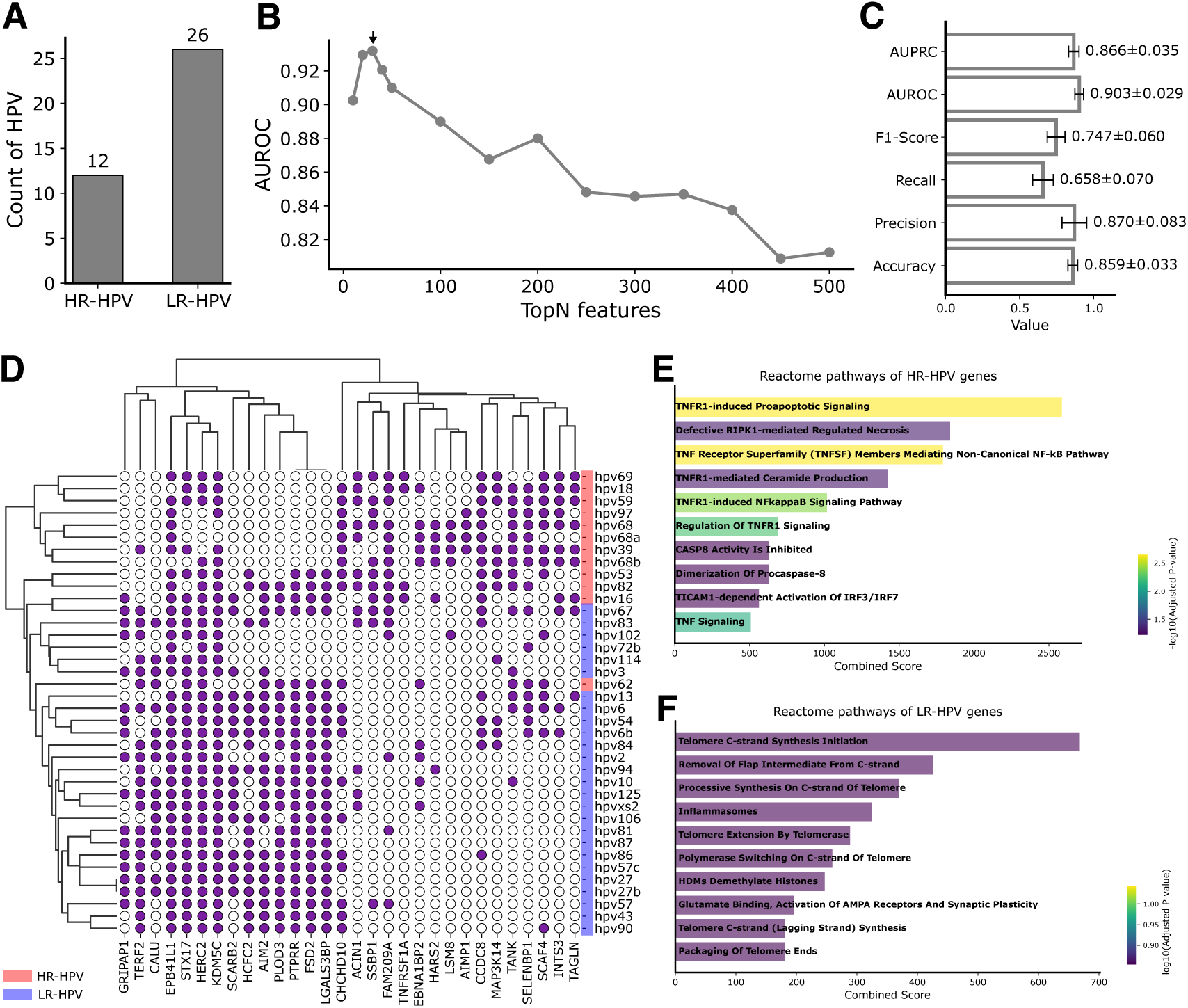
Classification of HPV based on virus-human PPIs. **A.** Number of high-risk (HR) and low-risk (LR) HPVs used for classification. **B.** AUROC across top N important features during feature selection. **C.** Performance metrics (accuracy, precision, recall, F1-Score, AUROC, and AUPRC) of combining the top 30 features with Random Forest classifier on the test set. **D.** Visualization of interactions between HPVs and the top 30 human genes. HR-HPVs are shown in red, and LR-HPVs in blue. Both HPVs and human genes were hierarchically clustered. **E-F.** Reactome enrichment analysis of human genes for HR-HPVs (**E**) and LR-HPVs (**F**).

## Discussion

The genetic makeup, defined as the genotype, produces proteins through gene expression and translation, and the proteins in turn influence the observable traits or phenotype of an organism. In diverse phenotype prediction tasks, genomic information is typically the first utilized. For example, Gokhman et al. predicted the direction of phenotypic difference (such as blood pressure, height, and metabolic rate) of human based on genomic data by statistical modeling[23]. Because proteins act as the functional executors of genetic information, lots of studies have extended phenotype prediction to the proteomic level. Zhao et al. predicted human disease phenotypes for early detection of diseases based on human protein sequences with a deep ensemble model[24]. Furthermore, as the regulation of biological pathways is orchestrated through interactions among biomolecules, particularly protein-protein interactions (PPIs), several studies have incorporated PPIs into phenotype prediction. For instance, Bi et al. predicted disease phenotype with a self-supervised learning approach that integrated PPIs with gene ontology information[25]. Collectively, these studies underscore the critical roles of genomes, proteomes, and PPIs in predicting phenotypic outcomes across biological systems.

Viral phenotype prediction presents unique challenge compared to cellular organisms as viruses rely entirely on host machinery for replication and survival. Viral phenotypes emerge not solely from viral genomes or proteomes but also from the dynamic interactions with host factors. Currently, numerous computational approaches have been developed for inferring viral phenotypes such as host range, infectivity, transmissibility and virulence, yet most rely solely on viral intrinsic information, particularly genomic or proteomic sequences. For example, Li et al. inferred multiple influenza virus phenotypes based on genomic sequences[8]; Peng et al. predicted the antigenic variation of influenza viruses based on HA protein sequences[26]. Some studies used the virological characteristics (such as genome type and tissue tropism) in inferring viral phenotypes. A few research have attempted to integrate host-related information for predicting vial phenotypes. For example, Li et al. combined spike protein of coronaviruses and virus receptor similarity among species to assess coronavirus cross-species transmission risk[27]. Taken together, existing methods for viral phenotype prediction mainly used the virus information, ignoring host determinants of virus phenotypes.

Virus-human PPIs are essential for viral entry, replication, immune evasion, and transmission. Thus, this study proposes that virus-human PPIs act as fundamental determinants of diverse viral phenotypes and therefore can be used for phenotype prediction. Unfortunately, the number of experimentally validated PPIs is far from complete. Our previous work established vhPPIpred[12] for large-scale prediction of virus-human PPIs, substantially expanding the interaction space available for downstream analyses. Building on these predicted interactions, we developed a PPI-based framework and systematically evaluated its performance across five viral phenotype prediction tasks. For human infectivity, our approach consistently outperformed sequence-based models, including those leveraging genome- or proteome-scale large language models. Additionally, compared with Mollentze’s method, the PPI-based framework also performed better. Similarly, for the remaining three phenotypes of human RNA viruses, PPI-based models showed superior predictive performance, with particularly strong improvements in inferring human transmissibility. The superiority of the PPI-based models can be attributed to their explicit incorporation of host factors which can capture the virus-host interaction patterns not included by virus sequence alone. For example, the human proteins used in predicting human infectivity were involved in virus infection and immune response, and the human proteins used in classifying HR-HPV and LR-HPV were extensively involved in virus infection and pathogenicity. Therefore, incorporating host determinants is essential for accurate viral phenotype prediction.

The PPI-based viral phenotype prediction framework established in this study are useful for assessing the risks of emerging viruses. It can be used to rapidly infer multiple phenotypes of newly discovered viruses including human infectivity, human transmissibility, transmission route, and tissue tropism, which are essential for risk evaluation of emerging viruses. Beyond these predictive applications, our approach holds potential for mechanistic studies. By identifying key human proteins related to viral phenotypes, our method can provide candidate host factors for experimental investigation, potentially revealing new therapeutic targets or biomarkers for infectious diseases.

Despite these advances, several avenues remain for improving the scope, accuracy, and generalizability of our framework. Firstly, the models for transmissibility, transmission route, and tissue tropism were trained exclusively on RNA viruses due to limited data. The relatively limited number of viruses with well-characterized phenotypes and potential uncertainty in categorical annotations due to incomplete or evolving epidemiological evidence may constrain model performance and generalizability. Expanding the dataset with more viruses especially DNA viruses and with more refined phenotype annotations should further improve model robustness. Secondly, as with other computationally inferred interaction datasets, predicted virus-human PPIs inevitably contain false positives and other prediction uncertainties. Although our computational validation supports the predictive value of PPI profiles for viral phenotype inference, it does not constitute experimental validation of individual predicted interactions or phenotypes. Prospective experimental and epidemiological validation of high-priority predictions will therefore be important. Thirdly, our current approach cannot resolve viral phenotypes at the strain level, representing an important direction for achieving higher-resolution predictions.

## Conclusion

This study introduces a perspective that virus-human protein-protein interactions (PPIs) serve as fundamental determinants of viral phenotypes. Based on this, we developed a computational framework to predict diverse viral phenotypes. Systematic evaluations demonstrated that PPI-based models consistently outperformed genome- and proteome-based approaches, exhibiting superior robustness and interpretability. Collectively, this work advances computational strategies for viral phenotype prediction and underscores the importance of incorporating virus-host interaction information into the early identification, prevention, and control of emerging viral threats.

## Materials and Methods

### Experimental-determined physical virus-human protein-protein interactions

Virus-human protein-protein interactions were obtained from the integrated PPI resource established in our previous study [12]. Briefly, virus-human PPIs were collected from eight public databases, including BioGRID (v4.4.208)[28], IntAct (v4.2.3.2)[29], VirusMentha (v1.0)[30], VirusMINT (v1.0)[31], VirHostNet (v3.0)[32], Viruses.STRING (v10.5)[33], HVIDB (v1.0)[34], and HVPPI (v1.0)[35], with data retrieved on May 16, 2022. Protein identifiers from different sources were mapped to UniProt accession numbers using bioDBnet (v2.1)[36]. To harmonize interaction confidence across databases, interaction scores were independently rescaled within each source using max-min normalization. The normalized datasets were subsequently merged, and redundant records representing the same interaction pair and interaction type were collapsed by retaining the entry with the highest normalized score, yielding 108,331 non-redundant virus-human interactions. To obtain a higher-confidence physical interaction set, we further retained records annotated with direct physical interaction keywords (“physical association”, “direct interaction”, “physical”, or “covalent binding”). When multiple records or interaction annotations were available for the same pair, the highest-scoring interaction was retained. Viral proteins shorter than 50 amino acids or longer than 1,000 amino acids were excluded to reduce potential contributions from short peptides and viral polyproteins. The resulting high-confidence dataset comprised 16,314 virus-human PPIs involving 938 viral proteins and 5,848 human proteins (Table S1).

### Viruses for human infectivity prediction

Human-infecting viruses were collected from four major sources: i) the Zhang’s study that curated 239 human-infecting viruses (retrieved on August 25, 2026); ii) the Mollentze’s dataset that contains 261 human-infecting viruses (retrieved on August 25, 2026); iii) Olival’s dataset containing 263 human-infecting viruses (retrieved on August 25, 2026); iv) the Virus-Host Database (v0.2.1), Viral Zone Database (v1.0), VIRION Database (v2.0), and the Human Virus Database (v1.0)[9,13,37–41]. Human-infecting viruses from published datasets (Zhang, Mollentze, and Olival) were merged directly, while those from databases were determined by searching keywords of “Human virus” or “Zoonotic” on August 25, 2026. All viruses from these sources were manually classified into three evidence levels: high-confidence viruses were defined as those reported to infect humans in at least one published study; middle-confidence viruses included those reported to infect human cell lines in the literature, or those with complete genomic sequences annotated with humans as the host in the NCBI GenBank or RefSeq databases; low-confidence viruses were defined as those annotated with humans as the host but with incomplete genomic sequences. In total, we obtained 331 high-confidence, 346 middle-confidence, and 792 low-confidence human infecting viruses. A total of 564 high-confidence non-human-infecting viruses were obtained from Mollentze’s study after removing those annotated as human-infecting in our dataset. For training and evaluating human infectivity models, we used the 677 high- and middle-confidence human-infecting viruses together with 564 non-human-infecting viruses (Table S2 and S3).

### RNA viruses for predicting virulence, transmissibility, transmission route, and tissue tropism

Viruses used for predicting virulence, transmissibility, transmission route, and tissue tropism were obtained from Brierley’s study (Table S5)[9,42]. The phenotype annotations and categorical definitions were adopted from this systematically curated dataset rather than newly defined in the present study. Brierley et al. compiled these epidemiological and biological traits through extensive literature curation, and we retained the original categorical definitions to ensure consistency with this established dataset. For virulence prediction, a total of 214 RNA viruses were collected and classified into two categories: low-level virulence (n=155) and high-level virulence (n=59). For transmissibility prediction, the same set of 214 RNA viruses were collected and classified into three categories by human-to-human transmission potential: low (no human-to-human transmission) (n=123), middle (limited human-to-human transmission) (n=31), and high (sustained human-to-human transmission) (n=60). For transmission route prediction, 195 RNA viruses with complete annotation records were included and classified into four categories: vector-borne (n=86), direct contact (n=54), respiratory (n=30), and faecal-oral (n=25). For tissue tropism prediction, 196 RNA viruses were selected and categorized into eight tropisms: viraemic (n=69), neural (n=31), systemic (n=28), gastrointestinal (n=25), respiratory (n=20), circulatory (n=12), vascular (n=7), and hepatic (n=4), with viruses in the circulatory, vascular, and hepatic categories merged into the “other” group (n=23).

### Viruses for HPV classification

We manually collected 38 HPV subtypes from Lasso’s study on September 2, 2024, resulting in a dataset of 12 high-risk HPVs associated with cervical cancer and 26 low-risk HPVs associated with benign warts[21].

### Prediction of virus-human PPIs

For predicting virus-human protein-protein interactions (PPIs), reference genomes of 2,285 viruses, including 1,241 used for human infectivity prediction, 214 for transmissibility, transmission route, virulence and tissue tropism prediction, 792 low-confidence human-infecting viruses, 38 for HPV classification, were downloaded from the NCBI Genome database using the NCBI Datasets command-line tool (v16.37.0) on August 25, 2026[43]. These genomes were translated into proteomes using Prodigal (v2.6.3) with the -p meta parameter[44].

Virus-human PPIs were subsequently predicted using our previously developed vhPPIpred [12] (https://github.com/ZyuanZhang/vhPPIpred.git). Briefly, vhPPIpred is an XGBoost-based model that integrates sequence embeddings and evolutionary information with network topology and viral molecular mimicry. Performance comparison suggested that vhPPIpred consistently outperformed nine state-of-the-art methods across our benchmark dataset and three independent datasets[12].

In this study, a specificity threshold of 0.99 was applied to vhPPIpred predictions, and the retained PPIs were used for downstream viral phenotype predictions. The predicted virus-human PPIs were encoded into a binary interaction matrix, with rows corresponded to virus species and columns to human proteins. Matrix entries were set to 1 if a predicted interaction existed between a virus and a human protein, and 0 otherwise.

### Construction of viral genome-based features

Viral genomes were embedded using two approaches: k-mer composition (k=3,4,5) and DNABERT-2 (https://github.com/MAGICS-LAB/DNABERT_2.git)[15]. These methods generated embedding vectors of length 64 (3-mer), 256 (4-mer), 1024 (5-mer), and 768 (DNABERT-2), respectively. For each virus species, all corresponding genomic embedding vectors were averaged to generate a single vector representing its genomic features. This procedure resulted in four genome-based features sets: genome-3mer, genome-4mer, genome-5mer, and genome-DNABERT-2.

### Construction of viral proteome-based features

Viral proteomes derived from genome translation were embedded using k-mer frequencies (k=1,2,3) and the ESM-2 protein language model (esm2_t30_150M_UR50D, v1.0.3)[16]. These methods yielded embedding vectors of length 20 (k=1), 400 (k=2), 8000 (k=3) and 640 (ESM-2), respectively. For each virus species, all proteomic embedding vectors were averaged to generate a vector representing its proteomic features. This procedure resulted in four proteome-based feature sets: proteome-1mer, proteome-2mer, proteome-3mer, and proteome-ESM-2.

### Evaluation metrics

Model performance was evaluated using standard classification metrics. For the binary classification task of human infectivity prediction, we computed Accuracy, Precision, Recall, F1-Score, the Area Under the Receiver Operating Characteristic Curve (AUROC), and the Area Under the Precision-Recall Curve (AUPRC).

The metrics were defined as follows:

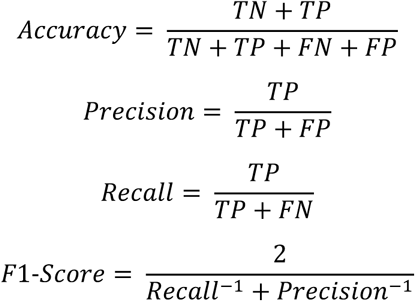

where TP, TN, FP and FN denote the number of true positives, true negatives, false positives and false negatives, respectively.

However, for multi-class prediction tasks (transmissibility, transmission route, and tissue tropism), we used the macro-averaged versions of these metrics, including Macro-Accuracy, Macro-Precision, Macro-Recall, Macro-F1-Score, Macro-AUROC, and Macro-AUPRC. Macro-averaging computes each metric independently for each class and then averages the results, giving equal weight to all classes.

All metrics were implemented using scikit-learn (v1.3.2)[45], including the functions accuracy_score, precision_score, recall_score, f1_score, roc_curve, precision_reall_curve and auc.

### Feature and algorithm selections for viral phenotype prediction

To identify the optimal combination of features and algorithms for predicting viral phenotypes (human infectivity, transmissibility, transmission route, and tissue tropism), three kinds of features (PPI-, genome-, and proteome-based features) were evaluated in combination with four machine-learning algorithms: Random Forest (RF), K-Nearest Neighbors (KNN), Support Vector Machine (SVM), eXtreme Gradient Boosting (XGBoost). Hyperparameters used for each algorithm were summarized in Table S7.

For selection of PPI-based features and algorithms, we ranked all human proteins using the SHapley Additive exPlanations (SHAP) scores computed with the shap package (v0.44.1)[46] in Python (v3.8.20)[47]. The top N (N=50, 100, … 2000) important human proteins were selected and combined with the four algorithms. For human infectivity prediction, viral samples were divided into training (70%), validation (15%), and test (15%) sets. Hyperparameters were tuned using 5-fold cross-validation on the training and validation sets via the grid search implemented by model_selection.GridSearchCV function of scikit-learn package., and each feature-algorithm combination was evaluated over 100 independent runs. The optimal feature-algorithm combination was chosen with the highest mean validation AUROC. This yielded PPI-Top1000-RF (named HI-PPI) for human infectivity, PPI-Top250-SVM (VL-PPI) for virulence, PPI-Top200-SVM (HT-PPI) for human transmissibility, PPI-Top350-SVM (named as TR-PPI) for transmission route, and PPI-Top1250-SVM (named as TT-PPI) for tissue tropism (Figure 2E, Figure S4A-D)

Genome-based features (genome-3mer, genome-4mer, genome-5mer, and genome-DNABERT-2) were evaluated with the four algorithms using the same cross-validation and repeated-run procedure. Based on mean validation performance (Accuracy, Precision, Recall, F1-Score, and AUROC), the model Genome-3mer-KNN (HI-Genome) was selected for human infectivity prediction (Figure S2A). For other tasks, Genome-4mer-KNN (HT-Genome), Genome-DNABERT-2-SVM (VL-Genome), Genome-4mer-SVM (TR-Genome), and Genome-DNABERT-2-RF (TT-Genome) achieved the highest mean validation AUROC (Figure S4E-H).

Proteome-based features (proteome-1mer, proteome-2mer, proteome-3mer, and proteome-ESM-2) were evaluated similarly. The optimal models were Proteome-ESM-2-RF (HI-Proteome) for human infectivity, Proteome-1mer-RF (HT-Proteome-TMS) for human transmissibility, Proteome-3mer-KNN (VL-Proteome) for virulence, Proteome-ESM-2-RF (TR-Proteome) for transmission route, and Proteome-ESM-2-RF (TT-Proteome) for tissue tropism (Figure S2B; Figure S4I-L).

For HPV subtype classification, human proteins were ranked by SHAP values, and the top N (N=10, 20, …, 500) proteins were evaluated using RF. The optimal number of features and the RF parameter n_estimators (5, 10, 15) were determined based on mean validation AUROC across 100 runs of 5-fold cross-validation. Finally, the top 30 human proteins were used for HPV classification. (Figure 5B).

### Comparison of PPI-based and Mollentze’s method

We adopted the data partitioning strategy of Mollentze et al. by using the publicly available code downloaded from https://github.com/Nardus/zoonotic_rank.git to generate the training, validation, and test sets. These identical data splits were used to enable a fair performance comparison between the PPI-based method and Mollentze’s approach. All feature selection, model training, and evaluation procedures of the PPI-based method were performed exclusively on the Mollentze’s dataset. The test AUROC of the PPI-based method was then directly compared with the AUROC reported for Mollentze’s method in the original publication (Figure S3A&B).

### Gene Ontology enrichment analysis of HI-PPI for human infectivity prediction

Human proteins included in the HI-PPI model were first converted to gene symbols using the mygene package (v3.2.2)[48–50] in Python (v3.8.20). These genes were then subjected to Gene Ontology biological process (GO-BP) enrichment analysis using the enrichGO function in the clusterProfiler (v4.12.6) package[51] in R (v4.4.1)[52]. Virus-related GO-BP terms were identified by filtering enriched terms using the keywords “viral” and “virus”. These terms were clustered based on semantic similarity computed with the pairwise_termsim function and visualized using the treeplot function of enrichplot package (v1.24.4)[53] in R. Similarly, GO-BP terms associated with immune responses identified using the keywords “immune”, “inflamma”, and “lymphocyte” were processed using the same clustering and visualization procedures.

### Reactome enrichment analysis of HPV-interacting human genes

After converting to gene symbols, the top 30 human proteins were hierarchically clustered using the clustermap function of seaborn (version 0.13.2)[54] in Python (v3.8.20), yielding two clusters corresponding to high-risk HPV-associated genes (HR-HPV group; 16 genes) and low-risk HPV-associated genes (LR-HPV group; 14 genes). Each gene group was then subjected to pathway enrichment analysis using the Reactome_2022 gene sets implemented by the enrichr function in the gseapy package (v1.1.9)[55]. Enrichment results were visualized using matplotlib (v3.7.5) in Python.

### Definition of risk score

The risk score used to rank potentially human-infecting viruses was calculated by combining the predicted scores of human infectivity, virulence, and sustained human-to-human transmissibility.

### Statistical analysis

All statistical analyses were performed using Python (v3.8.20). Data are presented as mean±SD (standard deviation), as shown in Figure 3J-M and Figure 5C. Comparison between two groups were conducted using two-sided t-test implemented with ttest_ind function of scipy package (v1.10.1)[56], including those shown in Figure 2F, Figure S1A&B and Figure S3B.

### Visualization of results

All figures in this study were generated using Python (3.8.20) in combination with Pandas (v2.0.3), NumPy (v1.24.4) and Matplotlib (v3.7.5), and using R (4.4.1) in combination with ggplot2 (v4.0.4), enirchplot (v1.28.4), UpSetR (v1.4.0) and ggsankey (https://github.com/davidsjoberg/ggsankey.git) [53,57–62].

## Data and materials availability

All data used in this study are provided in the supplemental materials. All analysis scripts and source code are publicly available at https://github.com/ZyuanZhang/vhPPIpredict-viral-phenotypes.git.

## Acknowledgements

This work was supported by the National Natural Science Foundation of China (32370700), Hunan Provincial Natural Science Foundation of China (2024JJ2015), and R&D Program of Guangzhou National Laboratory (GZNL2024A01002).

## Author contributions

Conceptualization: Z.Z., and Y.P. Methodology: Z.Z., and Y.F. Investigation: Z.Z., and Y.F. Visualization: Z.Z., and Y.F. Supervision: Z.Z., X.M., and Y.P. Writing—original draft: Z.Z., and Y.P. Writing—review & editing: Z.Z., X.G., X.M., and Y.P.

