## Supplementary material for "Virus-human protein-protein interactions predict viral phenotypes": Figure S1 to S5

**This file includes:**

Figure S1 to S5
References

**Figure S1. Comparison of the number of predicted virus-human PPIs used in human infectivity prediction. A.** Comparison of the number of predicted PPIs between human-infecting and non-human infecting viruses. **B.** Comparison of the number of unique human proteins between human-infecting and non-human-infecting viruses. **C.** Correlation between the length of viral genomic sequences and the number of predicted virus-human PPIs. **D.** Correlation between the number of viral proteins and the number of predicted virus-human PPIs.

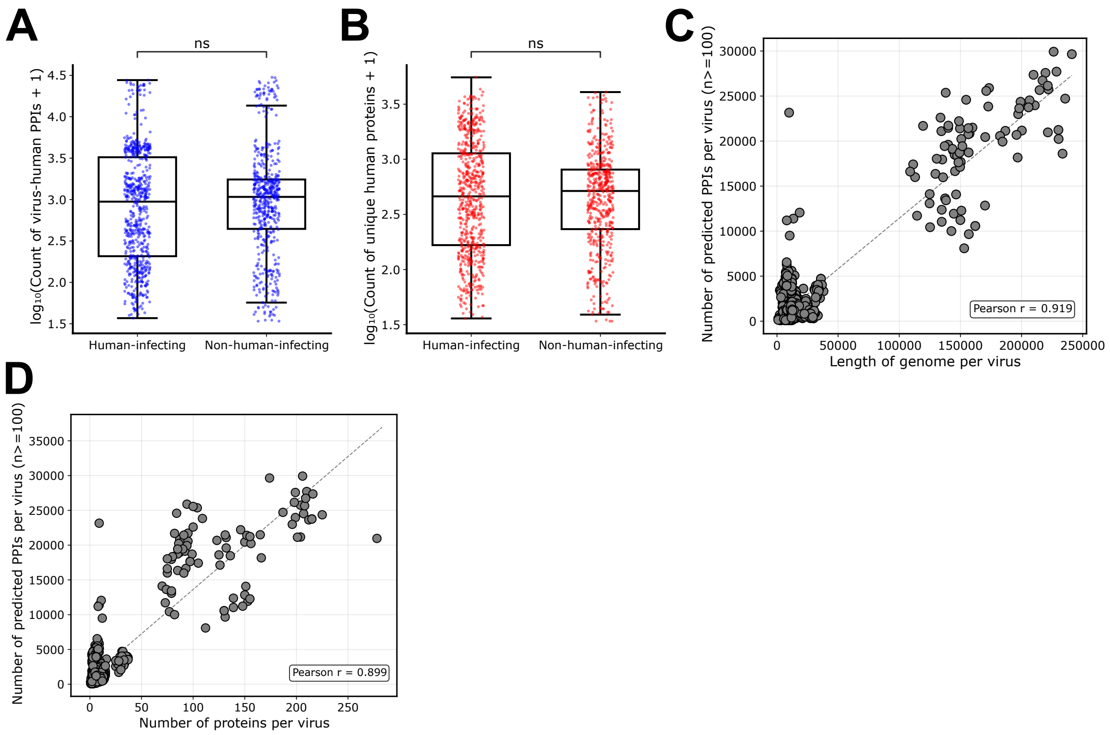

**Figure S2. Cross-validation performance of genome- and proteome-based models for predicting human infectivity. A.** Validation results combining genome-based features (Genome-3mer, 4mer, 5mer, and DNABERT2[1]) with four machine learning algorithms (KNN[2], RF[2], SVM[2], and XGBoost[3]). **B**. Validation results combining proteome-based features (Proteome-1mer, 2mer, 3mer, and ESM2[4]) with the same four algorithms. The optimal feature-algorithm combination was determined primarily based on the AUROC, with other metrics serving as secondary considerations. The colors of the heat map are filled based on the values normalized by min-max within each metric.

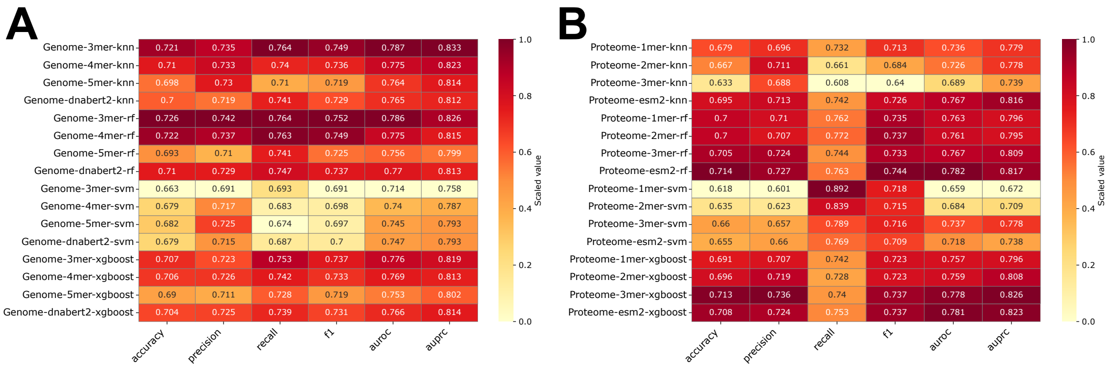

**Figure S3. Performance of Mollentze's and PPI-based model for predicting human infectivity on Mollentze's dataset. A.** Validation AUROC using the top N (N=50, 100, …, 2000) binary PPI features across the four algorithms. **B**. Test AUROC comparison of Mollentze's and PPI-based model on Mollentze's dataset[5].

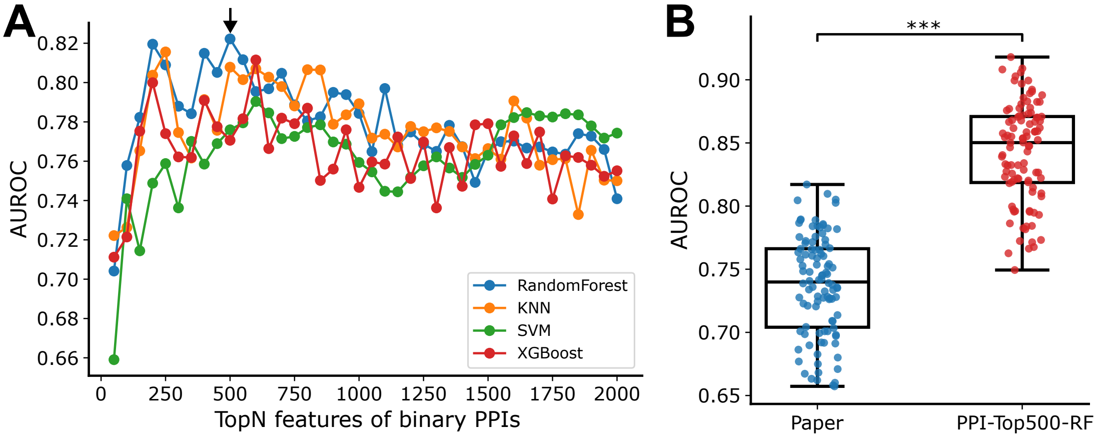

**Figure S4. Cross-validation results of predicting virulence (VL), human transmissibility (HT), transmission route (TR), and tissue tropism (TT) of viruses. A-D**. Validation AUROC using the top N (N=50, 100, …, 2000) binary PPI features across machine learning algorithms (KNN, RF, SVM, and XGBoost) on prediction of virulence (**A**), transmissibility (**B**), transmission route (**C**), and tissue tropism (**D**). **E-H.** Validation results combining Genome-based features (Genome-3mer, 4mer, 5mer, and DNABERT-2) with six machine-learning algorithms (KNN, SVM, XGBoost, and RF) for predicting virulence (**E**) transmissibility (**F**), transmission route (**G**), and tissue tropism (**H**). **I-L.** Validation results combining Proteome-based features (Proteome-1mer, 2mer, 3mer, and ESM-2) with the same six algorithms for predicting virulence (**I**), transmissibility (**J**), transmission route (**K**), and tissue tropism (**L**). Colors of heat map are filled based on the values normalized by min-max within each metric.

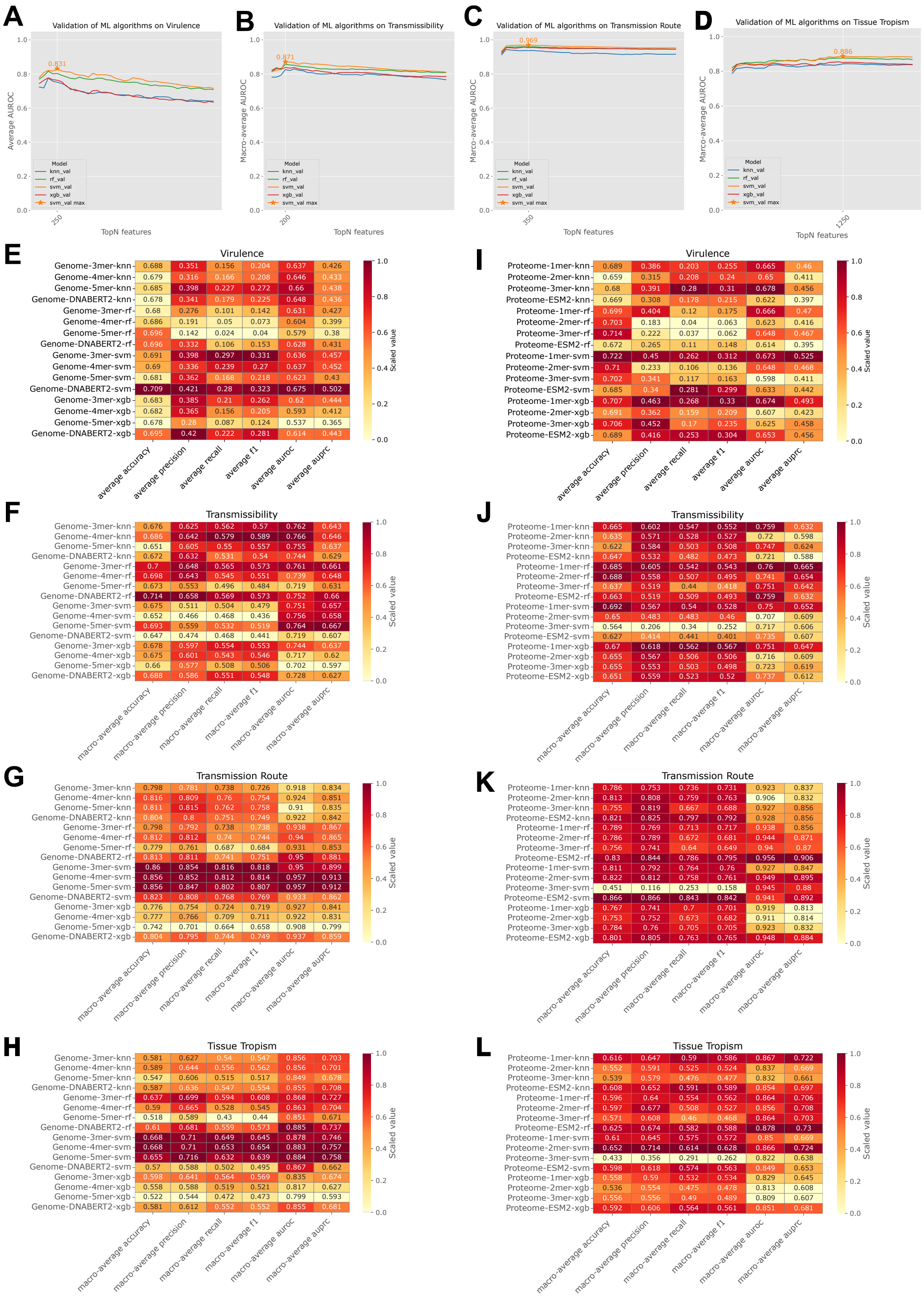
A
Validation of ML algorithms on Virulence
Average AUROC
TopN features
0.831
B
Validation of ML algorithms on Transmissibility
Macro-average AUROC
TopN features
0.871
C
Validation of ML algorithms on Transmission Route
Marco-average AUROC
TopN features
0.969
D
Validation of ML algorithms on Tissue Tropism
Marco-average AUROC
TopN features
0.886
Model
knn_val
rf_val
svm_val
xgb_val
svm_val max
E
Virulence
F
Transmissibility
G
Transmission Route
H
Tissue Tropism
I
Virulence
J
Transmissibility
K
Transmission Route
L
Tissue Tropism
Scaled value
average accuracy
average precision
average recall
average f1
average auroc
average auprc
macro-average accuracy
macro-average precision
macro-average recall
macro-average f1
macro-average auroc
macro-average auprc
Genome-3mer-knn
Genome-4mer-knn
Genome-5mer-knn
Genome-DNABERT2-knn
Genome-3mer-rf
Genome-4mer-rf
Genome-5mer-rf
Genome-DNABERT2-rf
Genome-3mer-svm
Genome-4mer-svm
Genome-5mer-svm
Genome-DNABERT2-svm
Genome-3mer-xgb
Genome-4mer-xgb
Genome-5mer-xgb
Genome-DNABERT2-xgb
Proteome-1mer-knn
Proteome-2mer-knn
Proteome-3mer-knn
Proteome-ESM2-knn
Proteome-1mer-rf
Proteome-2mer-rf
Proteome-3mer-rf
Proteome-ESM2-rf
Proteome-1mer-svm
Proteome-2mer-svm
Proteome-3mer-svm
Proteome-ESM2-svm
Proteome-1mer-xgb
Proteome-2mer-xgb
Proteome-3mer-xgb
Proteome-ESM2-xgb

**Figure S5. Human infectivity prediction for potentially human-infecting viruses using the HI-PPI.** Prediction results are generated by the HI-PPI model.

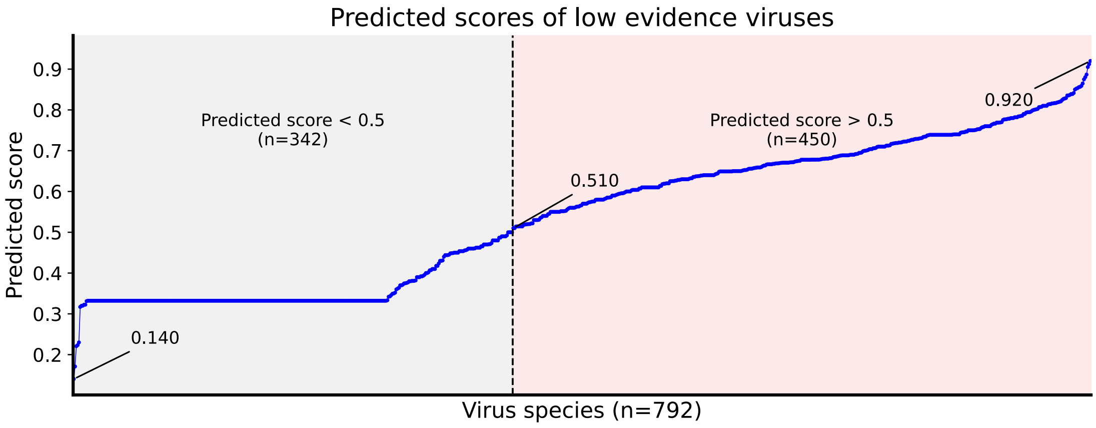
